# Genomes of all known members of a *Plasmodium* subgenus reveal paths to virulent human malaria

**DOI:** 10.1101/095679

**Authors:** Thomas D. Otto, Aude Gilabert, Thomas Crellen, Ulrike Böhme, Céline Arnathau, Mandy Sanders, Samuel Oyola, Alain Prince Okouga, Larson Boundenga, Eric Willaume, Barthélémy Ngoubangoye, Nancy Diamella Moukodoum, Christophe Paupy, Patrick Durand, Virginie Rougeron, Benjamin Ollomo, François Renaud, Chris Newbold, Matthew Berriman, Franck Prugnolle

## Abstract

*Plasmodium falciparum,* the most virulent agent of human malaria, shares a recent common ancestor with the gorilla parasite *P. praefalciparum.* Little is known about the other gorilla and chimpanzee-infecting species in the same (*Laverania*) subgenus as *P. falciparum* but none of them are capable of establishing repeated infection and transmission in humans. To elucidate underlying mechanisms and the evolutionary history of this subgenus, we have generated multiple genomes from all known *Laverania* species. The completeness of our dataset allows us to conclude that interspecific gene transfers as well as convergent evolution were important in the evolution of these species. Striking copy number and structural variations were observed within gene families and one, *stevor* shows a host specific sequence pattern. The complete genome sequence of the closest ancestor of *P. falciparum* enables us to estimate confidently for the first time the timing of the beginning of speciation to be 40,000-60,000 years ago followed by a population bottleneck around 4,000-6,000 years ago. Our data allow us also to search in detail for the features of *P. falciparum* that made it the only member of the *Laverania* able to infect and spread in humans.

## Main Text

The evolutionary history of *Plasmodium falciparum,* the most common and deadliest human malaria parasite, has been the subject of uncertainty and debate^1,2^. Recently it has become clear that *P. falciparum* is derived from a group of parasites infecting African Great Apes and known as the *Laverania* subgenus^2^. Until 2009, the only other species known in this subgenus was a parasite of chimpanzees known as *P. reichenowi,* for which only one isolate was available^3^. It is now clear that there are a total of at least seven species in Great Apes that naturally infect chimpanzees (P. *gaboni, P. billcollinsi* and *P. reichenowi),* gorillas (*P. praefalciparum, P. blacklocki* and *P. adleri*)^4,5^, or humans (*P. falciparum*) (Fig. 1a). Within this group, *P. falciparum* is the only parasite that has successfully adapted to humans after a transfer from gorillas and subsequently spread all over the world^2^.

**Figure 1.**
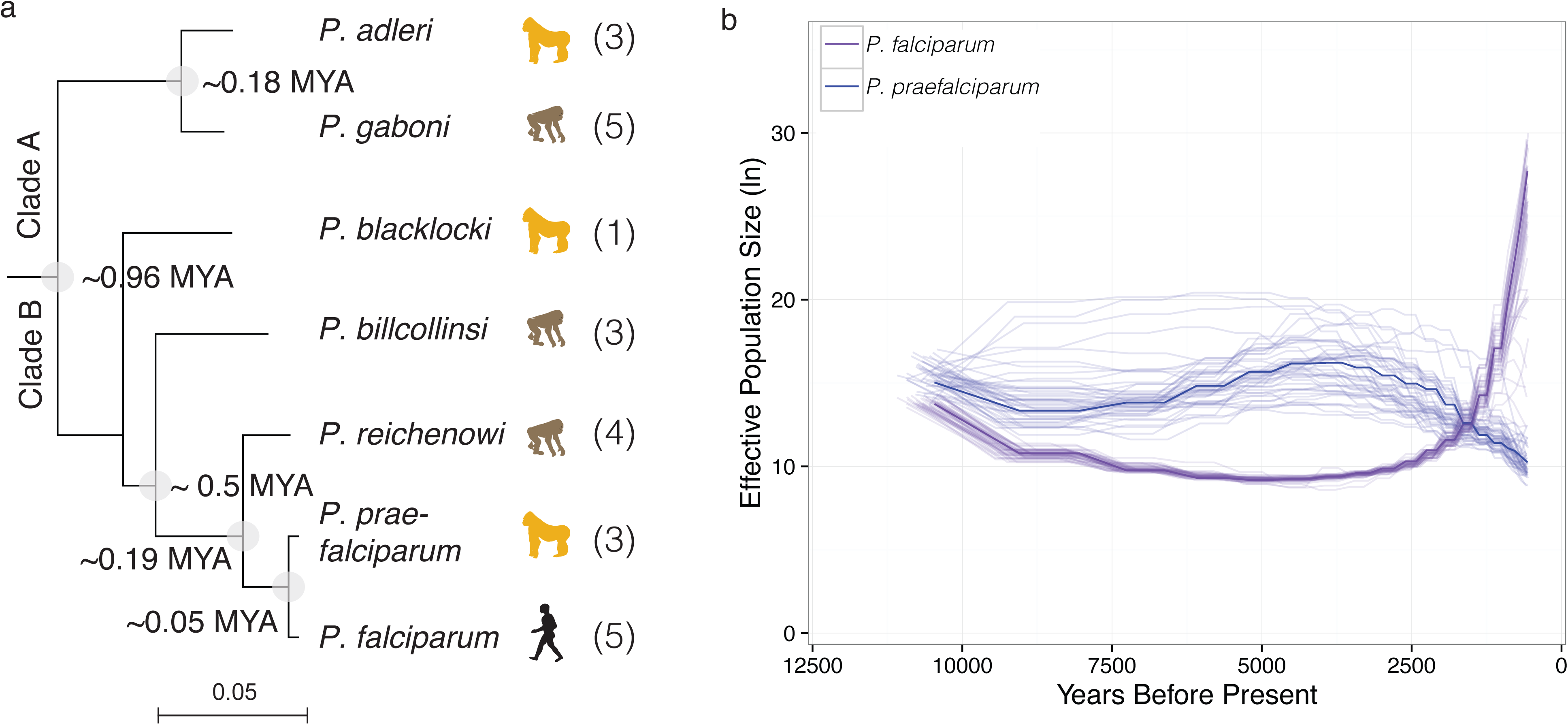
Overview of the dating of the evolution of the *Laverania.* (**a**) Maximum likelihood tree of the *Laverania* based on the “Lav12sp” set of orthologues. All bootstrap values are 100. Coalescence based estimates of the timing of speciation events are displayed on nodes (MYA - million years ago), based on intergenic and genic alignments. (**b**) Multiple sequentially Markovian coalescent estimates of the effective population size *(Ne)* in the *P. falciparum* and *P. praefalciparum* population. Assuming our estimate of the number of mitotic events per year, a bottleneck occurred in *P. falciparum* 4,000-6,000 years ago. The y-axis shows the natural logarithm (Ln) of *Ne.* Bootstrapping (pale lines) was performed by randomly resampling segregating sites from the input 50 times.

Over time there have been various estimates concerning the evolutionary history of *P. falciparum* with the speciation event having been estimated to be anywhere between 10,000 to 5.5 million years ago, the latter falsely based on the date of the chimpanzee–human split^6,7^. Others report a bottleneck less than 10,000 years ago^8^, but suggest a drop to a single progenitor parasite. The latter seems unlikely due to the presence of allelic dimorphisms that predate speciation events and therefore could not have both been transmitted if a new species were founded by a single individual infection. Also, the dating of the speciation cannot be accurately estimated without the genome sequence of *P. praefalciparum,* the closest living sister species to *P. falciparum.*

The absence of *in vitro* culture or a usable animal model has precluded obtaining sufficient DNA for full genome sequencing and has hindered investigation of the *Laverania.* So far just the full draft genome of *P. reichenowi* ^9^ and a nearly complete draft sequence of *P. gaboni*^6^ are available. These data together with additional PCR based approaches^10^ have provided important insights into the evolution of this subgenus, including the lateral gene transfer of the *rh5* locus, the early expansion of the FIKK gene family and the observation that the common ancestor also had *var* genes. Our data confirm and significantly extend these findings. However, the lack of whole genome information for the whole subgenus (particularly *P. praefalciparum*) has severely constrained the scope of subsequent analyses.

To investigate the evolutionary history of all known members of the *Laverania* subgenus and to address the question of why *P. falciparum* is the only extant species to have adapted successfully to humans, we have sequenced multiple genotypes of all known *Laverania* species.

### Genome sequencing from six *Laverania* species

Blood samples were taken during successive routine sanitary controls, from four gorillas and seven chimpanzees living in a sanctuary or quarantine facility prior to release (see Methods). A total of 15 blood samples were positive for ape malaria parasites by PCR. Despite low parasitemia in most animals, a combination of host DNA depletion, parasite cell sorting and amplification methods enabled sufficient parasite DNA templates to be obtained for short-read (Illumina) and long read (Pacific Bioscience) sequencing (Table 1). Mixed-species infections were frequent but resolved by utilising sequence data from single infections, resulting in 19 genotypes (Supplementary Table 1). The dominant genotype in each sample was assembled *de novo* (see Methods) using long read technology into a reference genome for six malaria parasite species: *P. praefalciparum, P. blacklocki, P. adleri, P. billcollinsi, P. gaboni* and *P. reichenowi.* The assemblies comprised 44-97 scaffolds (Table 1), with large contigs containing the subtelomeric regions and internal gene clusters that house multigene families known in *P. falciparum* and *P. reichenowi* to be involved in virulence and host-pathogen interactions. The high quality of the assemblies compared to those obtained previously is illustrated by the good representation of multi-gene family members (Supplementary Table 2) and the large number of one-to-one orthologues obtained between the different reference genomes (4,350 among the seven species and 4,826 between *P. falciparum, P. praefalciparum* and *P. reichenowi).* Two to four additional genomes were obtained for each species except for *P. blacklocki* (Table 1, Supplementary Table 1).

**Table 1:**
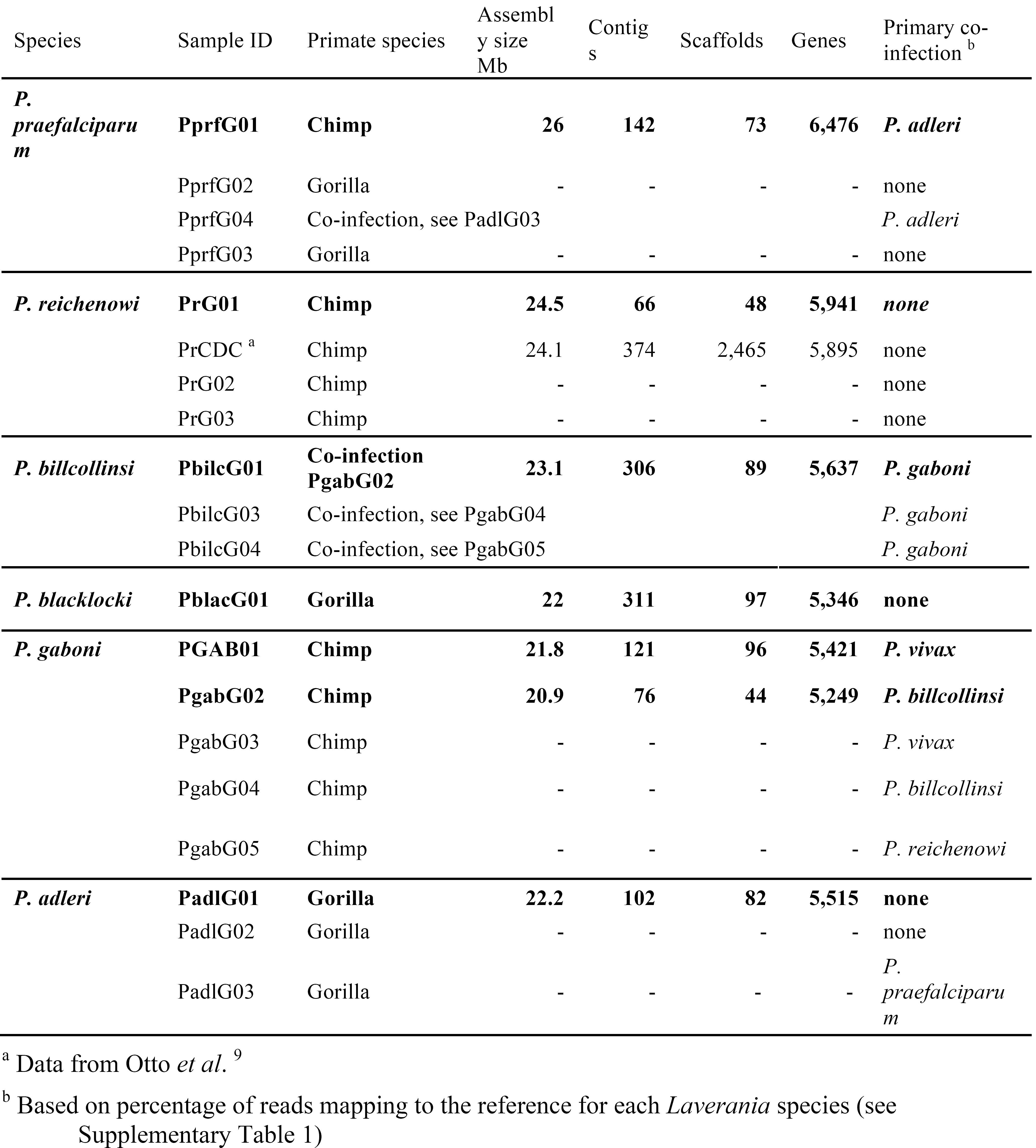
Overview of all *Laverania* samples used in study. All samples generated for this study, except PrCDC, were sequenced using Illumina technology with a range of read lengths from 100–250 bp. For isolates in bold, assemblies were produced using long-reads from Single-Molecule Real-Time sequencing (Pacific Biosciences) and the total assembled size, number of contigs, scaffolds and predicted genes are shown. Fold coverage is reported based on mapping reads to the P. falciparum 3D7 reference genome (v3.1). *P. billcollinsi* sequences were obtained from *P. gaboni* co-infections, of which some also harboured *P. reichenowi species*.

### Speciation history in the *Laverania* sub-genus

Conservation of gene content and synteny is striking between these complete genomes and enabled us to reconstruct with confidence the relationships between different *Laverania* species, to compare their relative genetic diversity (Fig 2a, Supplementary Fig. 1) and to estimate the age of the different speciation events that led to the extant species. The latter has been problematic in the past due to the lack of both complete genome data and accurate estimates of mutation rate and generation time. Using the most divergent estimates of generation time and measured mutation rates from the *P. falciparum* literature, we found the data converge to 0.9-1.5 mutations per year per genome (Supplementary Note 1). We observed a similar substitution rate *in vivo* by examining existing sequence data for five geographically diverse isolates, covering a 200-kb region surrounding the PfCRT gene that is relatively conserved due to a selective sweep resulting from chloroquine use (Supplementary Note 1; Supplementary Figure 2). The fact that these two figures are similar suggests that the *in vitro* mutation rate may have been underestimated since many mutations will be lost by genetic drift. Since no data is available for the other species we have assumed hereon that these values generalise across the subgenus. From our Bayesian whole-genome estimates, the ancestor of all current day parasites of this subgenus existed 0.7–1.2 million years ago, a time at which the subgenus divided into two main clades, A (P. *adleri* and *P. gaboni)* and B that includes the remaining species (Fig. 1a). Our range of values is far more recent than previous estimates^3,11^. Following the Clade A/B subdivision, several speciation events occurred leading either to new chimpanzee or gorilla parasites. Interestingly, the divergence between *P. adleri* and *P. gaboni* in one lineage and *P. reichenowi* and the ancestor of *P. praefalciparum/P. falciparum* in the other lineage occurred at approximately the same time (140–230 thousand years ago; Fig. 1a, Table 2). Based on our coalescence estimates, *P. falciparum* begun to emerge in humans from *P. praefalciparum* around 40–60 thousand years ago (Fig. 1a), significantly later than the evolution of the first modern humans and their spread throughout Africa^12^. Our analysis also indicates significant gene flow between these two parasite species after their initial divergence (Table 2).

**Table 2:** Estimated dates of speciation (time to coalescence), effective population size (Ne) and migration rates for present and ancestral *Laverania* populations. Estimated dates of speciation (time to coalescence), effective population size (Ne) and migration rates for Laverania populations, including inferred ancestors. Values inferred using GPhoCS coalescence model. The results are presented from sequence alignments of protein coding (CDS) (genic) regions as well as apparent intergenic regions inferred in the presence or absence of 0.5 kb flanking (UTR) sequences. The coalescence model parameters have been scaled based on an assumption of 401–681 mitotic events per year (Supplementary Note 1). The number in parentheses is the range based on the prediction (95% confidence interval). Values with (*) are linear estimates, as good alignments could not be obtained between those species due to the low GC content. The migration rate refers to the probability of a parasite from the source lineage migrating into the target over the time period. The data from these estimates are presented in Fig. 1.

| Population | Status | Coalescence, years ($\times 10^5$) | | | Effective Population size, $N_e$ ($\times 10^5$) | | | Migration target | Probability of Parasite Migration | | |
| --- | --- | --- | --- | --- | --- | --- | --- | --- | --- | --- | --- |
|  |  | CDS | Intergenic | Intergenic no UTR | CDS | Intergenic | Intergenic no UTR |  | CDS | Intergenic | Intergenic no UTR |
| *P. falciparum* | Present | NA | NA | NA | 1.9 - 3.2<br>(1.8 - 3.4) | 0.4 - 0.7<br>(0 - 2.1) | 0.4 - 0.7<br>(0 - 2.1) | NA | 0 | 0 | 0 |
| *P. praefalciparum* | Present | NA | NA | NA | 8.8 - 14.8<br>(8.1 - 16) | 12.2 - 20.7<br>(11.6 - 21.8) | 10 - 17<br>(9.2 - 18.4) | Pf | 0.02<br>(0.0 - 0.03) | 0.14<br>(0.09 - 0.19) | 0.16<br>(0.08 - 0.27) |
| *P. reichenowi* | Present | NA | NA | NA | 11.3 - 19.2<br>(3.7 - 19.9) | 15.9 - 26.9<br>(15.4 - 27.7) | 12.3 - 20.7<br>(11.7 - 21.8) | NA | 0 | 0 | 0 |
| *P. billcollinsi* | Present | NA | NA | NA | 9.3 - 15.8<br>(8.9 - 16.5) | - | - | NA | 0 | - | - |
| *P. gaboni* | Present | NA | NA | NA | 18.1 - 30.6<br>(17.6 - 31.6) | - | - | NA | 0 | - | - |
| *P. adleri* | Present | NA | NA | NA | 17.7 - 29.9<br>(16.9 - 31.2) | - | - | NA | 0 | - | - |
| *Pf - Pprf* | Ancestral | 0.2 - 0.3<br>(0.1 - 0.3) | 0.5 - 0.8<br>(0.4 - 0.8) | 0.4 - 0.6<br>(0.3 - 0.7) | 6.5 - 11.1<br>(6 - 11.9) | 16.5 - 27.9<br>(15.4 - 29.7) | 12.4 - 21<br>(11.1 - 23.2) | NA | 0 | 0 | 0 |
| *Pf - Pprf - Pr* | Ancestral | 0.9 - 1.5<br>(0.8 - 1.5) | 1.6 - 2.7<br>(1.5 - 2.7) | 1.4 - 2.3<br>(1.3 - 2.4) | 29.7 - 50.3<br>(27.7 - 53.7) | 68.6 - 116.2<br>(65.4 - 121.6) | 42.3 - 71.5<br>(39.1 - 77.1) | NA | 0 | 0 | 0 |
| *Pf - Pprf - Pr - Pbilc* | Ancestral | 2.3 - 4<br>(2.3 - 4) | 4.2 - 7.1* | 3.7 - 6.2* | 65.8 - 111.3<br>(62 - 118.5) | - | - | Pgab - Padl | 0.605<br>(0.45 - 0.77) | - | - |
| *Pgab - Padl* | Ancestral | 0.9 - 1.4<br>(0.8 - 1.5) | 1.5 - 2.6* | 1.3 - 2.2* | 28.3 - 47.9<br>(26.7 - 50.7) | - | - | NA | 0 | - | - |
| *All species* | Ancestral | 4.5 - 7.6<br>(4.2 - 8) | 8.1 - 13.6* | 7 - 11.8* | 98.9 - 167.4<br>(93.2 - 178.4) | - | - | NA | 0 | - | - |

*P. falciparum* has strikingly low diversity (*π* =0.0004), compared with the other *Laverania* species (0.002–0.0049) (Supplementary Fig. 1). It has been proposed that *P. falciparum* arose from a single transfer of *P. praefalciparum* into humans^6^ and based in part on the paucity of neutral SNPs within the genome, that *P. falciparum* emerged from a bottleneck of a single parasite around 10,000 years ago, after agriculture was established^6,8^. In light of our results, we estimate that the *P. falciparum* population declined around 11,000 years ago and reached a minimum about 5,000 years ago (Fig. 1b) with an effective population size (*Ne*) of around 3,000 (Supplementary Note 1; generally the census number of parasites is higher than Ne^13^). It is important to note that the methods we used to estimate *Ne* do not rely on the Fisher-Wright model, that could give problems as previously reported^14^. The hypothesis of a single progenitor is also inconsistent with the observation of several ancient gene dimorphisms that have been observed in *P. falciparum.* A previous analysis using *P. reichenowi* and *P. gaboni* sequence data, provided some evidence that different dimorphic loci diverged at different points in the tree^15^. Looking at each of these *P. falciparum* loci across the *Laverania,* we found different patterns of evolution at the *msp1, var1csa,* and *msp3* loci (Supplementary Fig. 3a). Most strikingly, a mutually exclusive dimorphism (described as MAD20/K1^16^) in the central 70% of the *msp1* sequence, clearly pre-dates the *P. falciparum–P*. *praefalciparum* common ancestor and dimorphism in *var1csa* (an unusual *var* gene of unknown function that is transcribed late in the asexual cycle) occurred before the split with *P. reichenowi*.

In contrast, the gene *eba-175* that encodes a parasite surface ligand involved in red blood cell invasion contains a dimorphism that arose after the emergence of *P. falciparum* (Supplementary Fig. 3b). The time to the most recent common ancestor of *eba-175* has been estimated as 130–140 thousand years in an analysis^17^ that assumed *P. falciparum* and *P. reichenowi* diverged 6 million years ago. However, based on our new estimate for *P. falciparum–P. reichenowi* divergence, we recalibrated their estimate of the most recent common ancestor of the *eba-175* alleles to be around 4,000 years ago, which is in good agreement with our divergence time for *P. falciparum* (Supplementary Note 1). The recent dimorphism cannot however explain the observation of an ancient dimorphism near the human and ape loci for glycophorin^18^ – an EBA-175 binding protein. The formation and maintenance of all of these dimorphic loci has therefore been shaped by different balancing selection pressures over time.

### *P. falciparum*-specific evolution

During its move away from gorillas, *P. falciparum* had to adapt to new environmental conditions, namely a new vertebrate host (human) and new vector species (e.g. *Anopheles gambiae*)^19^. To infer *P. falciparum* specific adaptive changes, we considered the *P. falciparum* /

*P. praefalciparum /P. reichenowi* genome trio and then applied two lineage based tests to find positive selection that occurred in the *P. falciparum* branch (see methods). The two tests identified 172 genes (out of 4,826) with signatures of positive selection in the human parasite species only (Supplementary Table 3). Two genes (*rop14* and PF3D7_0609900) were significant in both tests. Among the 172 genes, almost half (n=82) encoded proteins of unknown function. Analysis of those with functional annotation indicated that genes involved in pathogenesis, entry into host, actin movement and organization and in drug response were significantly over-represented. Other genes, expressed in different stages of the *P. falciparum* life cycle (e.g. *sera4* and *emp3,* involved during the erythrocytic stages; *trsp* and lisp1, involved in the hepatic stages; and *plp4, CelTOS* and *Cap380,* involved in the mosquito stages) also showed a significant signal of adaptive evolution (Supplementary Table 3).

### Evolution through introgression, gene transfer and convergence

Frequent mixed species infections in apes and mosquitoes^19^ provide clear opportunities for interspecific gene flow between these parasites. A recent study^6^ reported a gene transfer event between *P. adleri* and the ancestor of *P. falciparum* and *P. praefalciparum* of a region on chromosome 4 including key genes involved in erythrocyte invasion (*rhr5* and *cyrpA*). Because such events preserve the phylogenetic history of the genes involved, we systematically examined the evidence for introgression or gene transfer events across the complete subgenus by testing the congruence of each gene tree to the species tree for genes with one-to-one orthologues. Beyond the region that includes *rh5* (Fig 2b, Supplementary Fig. 4a), few signals of gene flow between parasites infecting the same host species were obtained (n=11) suggesting that these events were rare or usually strongly deleterious (Supplementary Fig. 5).

The *Laverania* subgenus evolved to infect chimpanzees and gorillas but, on a genome-wide scale, the convergent evolution of host-specific traits has not left a signature (Supplementary Note 2). We therefore examined each CDS independently and were able to identify genes with differences fixed within specific hosts, falling into three categories: 53 in chimpanzee-infective parasites, 49 in gorilla-infective and 12 with fixed traits in both host species (Fig. 2; Supplementary Table 4a). For at least 67 genes, these differences were unlikely to have arisen by chance (p <0.05) and GO term enrichment analysis revealed that several of these genes are involved in host invasion and pathogenesis (Supplementary Table 4b) including *rh5* (which has a signal for convergent evolution even when the introgressed tree topology is taken into consideration; Supplementary Fig. 4b). *Rh5* is the only gene identified in *P. falciparum* that is essential for erythrocyte recognition during invasion, via binding to Basigin. *P. falciparum rh5* cannot bind to gorilla Basigin and binds poorly to the chimpanzee protein^20^. We notice that one of the convergent sites is known to be a binding site for the host receptor Basigin^21^ (Supplementary Fig. 4b). The gene *eba-165* encodes a member of the erythrocyte binding like (EBL) super family of proteins that are involved in erythrocyte invasion. Although *eba-165* is a pseudogene in *P. falciparum*^22^, it is not a pseudogene in the other *Laverania* species and may therefore be involved in erythrocyte invasion, like other EBL members. The protein has three convergent sites in gorillas. One falls inside the F2 region, a domain involved in the interactions with erythrocyte receptors. The role of this protein and of these convergent sites in the invasion of gorilla red cells remains to be determined. Finally, genes involved in gamete fertility (the 6-cysteine protein P230) or implicated in *Plasmodium* invasion of erythrocytes (doc2^23^) also displayed signals of convergent evolution. Twelve parasite coding sequences had fixed differences at the same amino-acid position in chimpanzees and gorillas. Of these P230 was the only one found with a position that was different and fixed across all three host species. P230 is involved in gamete development and trans-specific reproductive barriers^24^, possibly through enabling male gametes to bind to erythrocytes prior to exflagellation^25^. Host-specific residues observed in P230 might affect the efficiency of the binding to the erythrocyte receptors and result from co-evolution between the parasite molecule and the host receptor.

**Figure 2.**
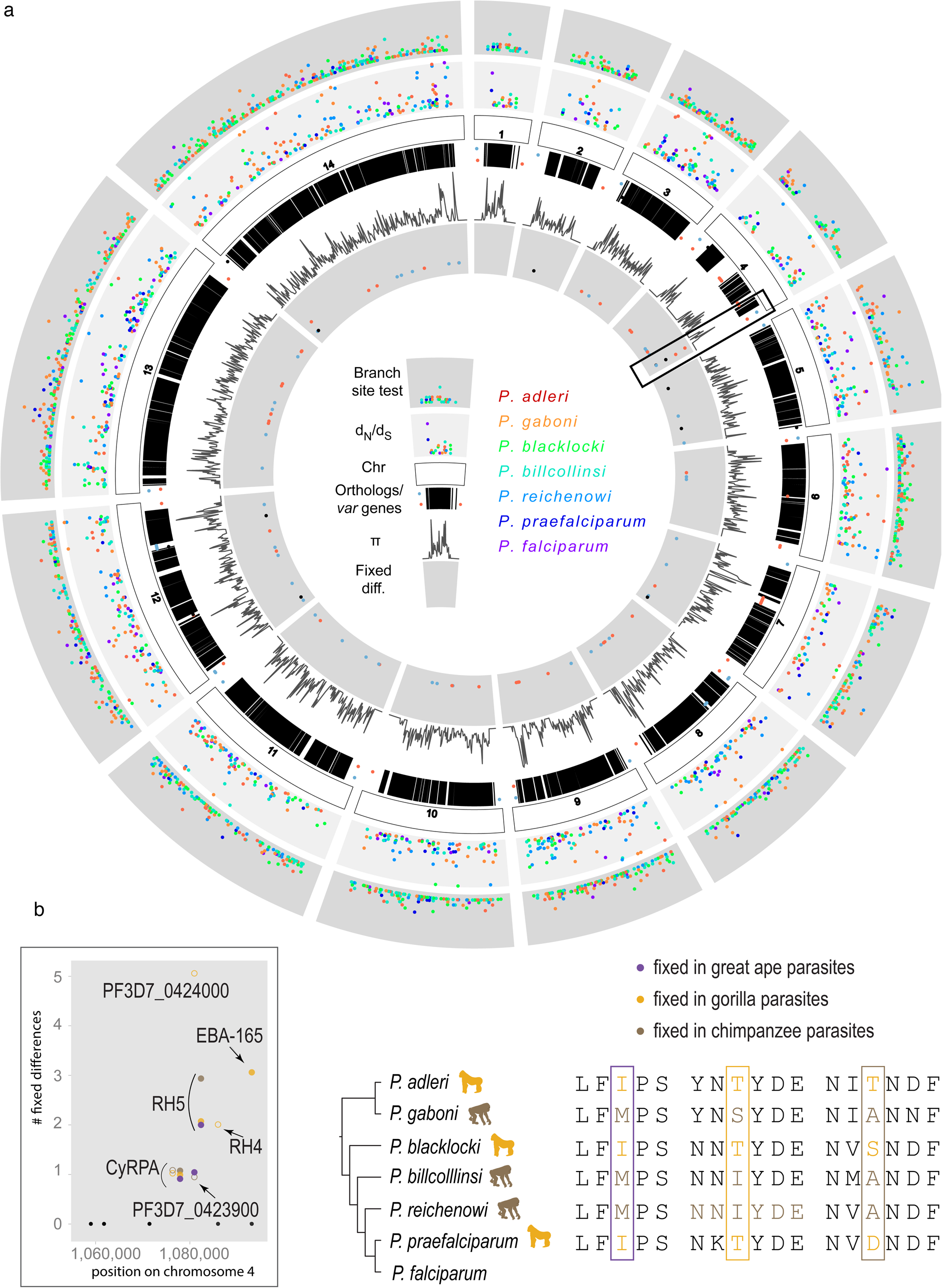
Overview of the analyses of core genes over all *Laverania* genomes. (a) Summary of evolution of core genes. From outer to inner track: scatterplot of branch site test for each genome (see Supplementary Table 3 for *P. falciparum* data); per-species *d*_N_/*d*_S_ values (0.5 < *d*_N_/*d*_S_ < 2); orthologues represented by vertical black lines under the chromosome track represent, with dots representing *P. falciparum* 3D7 *var* genes on the forward (blue) or reverse strands (red), or *var* pseudogenes (black); average of the relative polymorphism (n) across species, with the underlying n for each species calculated from multiple strains (“Lav25st” dataset) and normalized by the average for that species; signatures of convergent evolution based on host-specific fixed differences analysis with the chromosome 4 region that includes the *Rh5* locus highlighted (black box). (b) Magnified view of the *Rh5* region that is enriched with host specific fixed differences. Convergent evolution analysis was performed using orthologues conserved across the *Laverania*. Filled circles represent the subset of differences that were fixed within all the isolates available (“Lav15st” set) and for which we could reject neutral evolution (for the gene list see Supplementary Table 4).

### Subtelomeric gene families

To date, the only in-depth data on the subtelomeric gene families of the *Laverania* have come from *P. reichenowi* and *P. falciparum.* As a result of using long read sequencing, these families are well represented in our assemblies (Supplementary tables 2 and 5A). We provide for the first time a comprehensive picture of the evolution of these important families.

Most gene families were likely present in the ancestor of all Laverania. The same general pattern of one-to-one orthology throughout the subgenus indicates that many underwent gene duplication early (e.g. FIKK) or prior (e.g. ETRAMP, PHIST and SURFIN) to the development of a distinct Laverania lineage. Only a subset displayed contractions or expansions between specific Laverania species (Fig. 3 and Supplementary Table 5a, Supplementary Figure 7). For these latter families, Clade A and most species of Clade B clearly differ in their composition. *P. blacklocki* (Clade B) is intermediate in its composition. Some gene families, like the group of exported proteins *hyp4, hyp5, mc-2tm* and *EPF1,* have expanded only in *P. praefalciparum* and *P. falciparum* (and even more in *P. falciparum* for *hyp4* and *hyp5*). Since all four are components of Maurer’s clefts, an organelle involved in protein export^26^, some evolution of function in this organelle may have been an important precursor to human infection. The family of acyl-CoA synthetase genes, reported to be expanded and diversified in *P. falciparum*^27^ is in fact expanded across the *Laverania* and has four fewer copies in *P. falciparum* (Supplementary Fig. 6). Other genes that show clade or group specific expansion include DBLmsp, glycophorin binding protein and CLAG (Supplementary Fig. 7).

**Figure 3.**
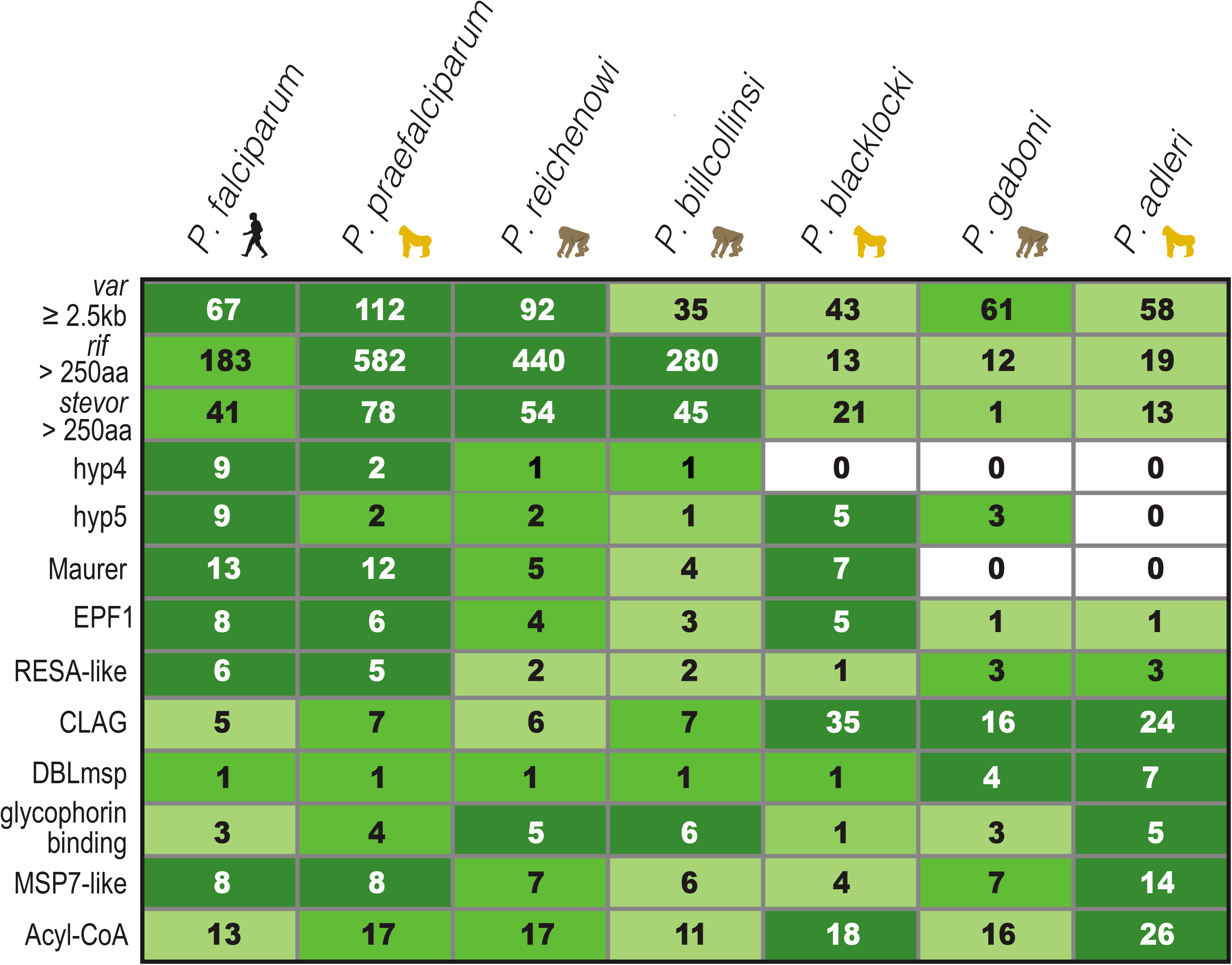
Gene families in the *Laverania.* Distribution of major multigene families including *var* and those that show significant copy number variation among lineages. Data from *P. praefalciparum* include the subtelomeric gene families from the two infecting genotypes. Assembly of *P. billcollinsi* is incomplete in the subtelomeres.

One striking inter-clade difference concerns the largest gene family that is likely common to all other malaria species: the *Plasmodium* interspersed repeat family *(pir,* which includes the *rif* and *stevor* families in *P. falciparum)* (Fig. 3, 4). This family has been proposed to be involved in important functions such as antigenic variation, immune evasion, signalling, trafficking, red cell rigidity and adhesion^28^ and yet has expanded only in Clade B, after the *P. blacklocki* split (Fig 3). The *rif* genes comprise a small conserved group and a much larger group of more diverse members that contains just 13 genes from Clade A species and at least 180 members per Clade B species (Fig. 4). There is however no evidence for host-specific adaptation in these sequences.

**Figure 4:**
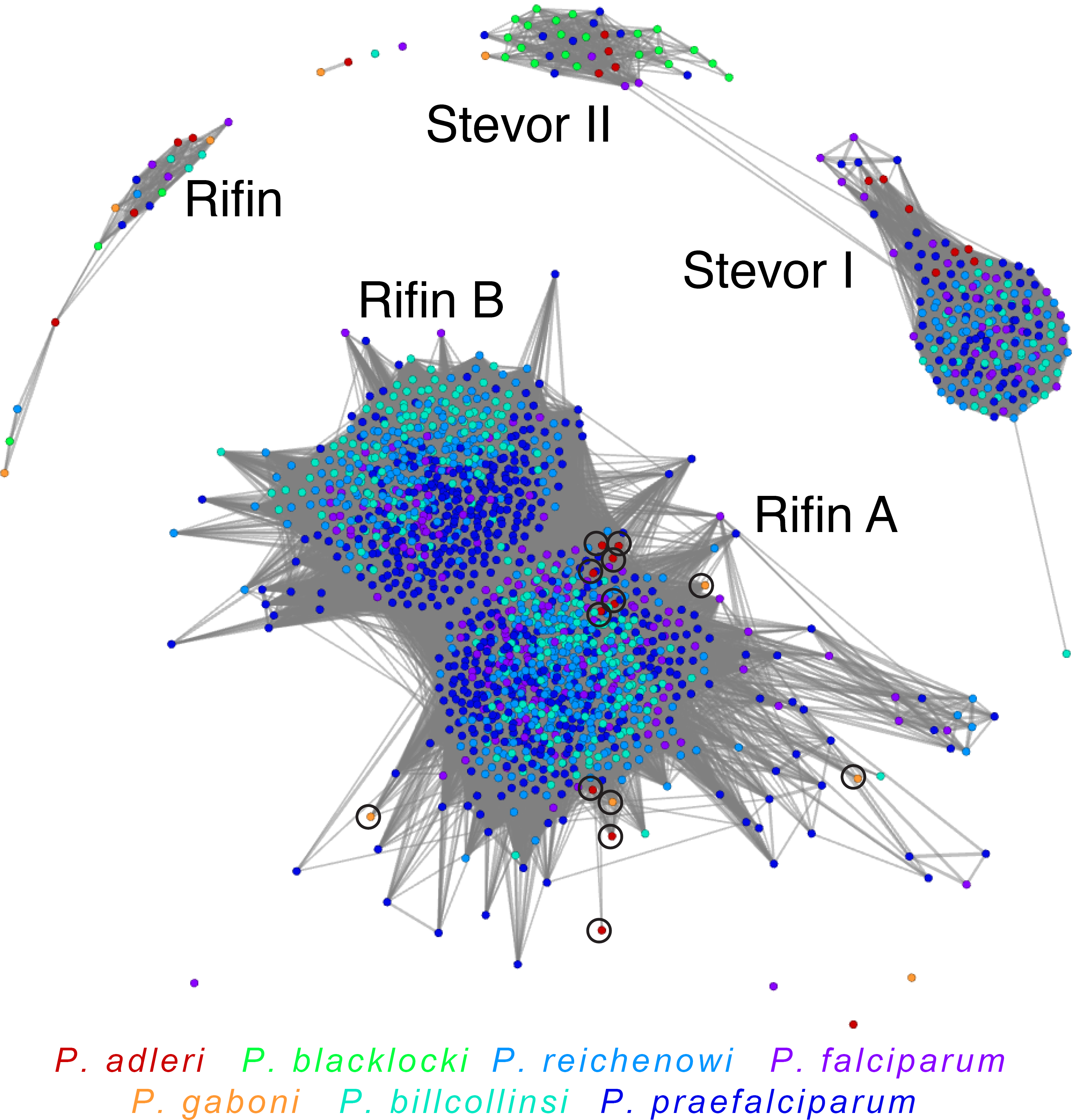
Clustering of Pir (Rifin and Stevor) proteins families. Graphical representation of similarity between all *pir* proteins > 250aa, coloured by species. A BLAST cut-off of 45% global identity was used (see methods). More connected genes are more similar. Black circles highlight Clade A rifin proteins that cluster with Clade B rifin proteins.

In contrast, a subset of *stevor* genes showed strong host-specific sequence diversification (Fig 4 and Supplementary Fig. 8). Based on full-length alignments, there is a deep phylogenetic split between *stevor* genes but when only short conserved protein motifs are considered, a group of Stevor proteins *(stevor II,* Fig 4a) forms a cluster comprising almost entirely of members from gorilla-infecting species. Since *stevor* genes are known to be involved in host-parasite interactions (such as binding to host glycophorin C in *P. falciparum^29^),* this host specific sequence may reflect sequence differences in host-specific factors in gorillas.

### Evolution of *var* genes

The *var* genes, crucial mediators of pathogenesis and the establishment of chronic infection through cytoadherence and immune evasion, are the best studied *P. falciparum* multi-gene family and unique to the *Laverania*^30^. They are two-exon genes and their products have three types of major domain; exon 1 encodes Duffy Binding like (DBL) and Cysteine Rich Interdomain Regions (CIDR) and exon 2 encodes Acidic Terminal Sequence (ATS)^31^. Similar to *P. falciparum,* our data are consistent with all *Laverania* species having *var* genes (Fig. 3) that retain a two-exon structure and are organized into subtelomeric or internal *var* gene clusters. There are however three notable features of the evolution of this family within the sub-genus.

First, there is a deep division in how the repertoire is organised between the major clades. The *var* genes of Clade B parasites, with the exception of *P. blacklocki,* resemble those of *P. falciparum* in terms of genomic organisation, domain types and numbers (Fig 5, Supplementary Table 6). In contrast, the repertoires of Clade A parasites and *P. blacklocki* (treated as one group hereafter in this section) differ in their domain composition, contain a novel CIDR-like domain (CIDRn, Fig 5a, Supplementary Fig. 9) and have lower sequence diversity per domain but cluster into more sub-groups than Clade B domains (Fig 5b, Supplementary Fig. 10). The paucity of domains similar to those in *P. falciparum* (such as CIDRα) that are involved in cytoadherence to some specific and common host receptors, means that if endothelial cytoadherence was important in Clade A, some alternative receptors must have been utilised.

Second, in total there are 10 internal *var* gene clusters (confirmed by contiguous sequence data) but 8 are oppositely oriented between the two clades (Supplementary Fig. 11, Supplementary Table 7). Clade B parasites also show a much greater number of associated GC-rich RNAs of unknown Function (RUF) elements than Clade A (Supplementary Table 7).

Third, the ATS domains cluster tightly within Clade A. Within Clade B there is clear evidence of species-specific diversification, except in *P. praefalciparum* and *P. falciparum* reflecting their recent speciation. There is one intact ATS from *P. falciparum* as well as several pseudogenes that cluster with Clade A (Fig 5c). Moreover, of seven internal *var* arrays (Supplementary Fig. 11) in *P. falciparum,* containing a functional *var* gene, five terminate with one of these pseudogenes (on the opposite DNA strand) suggesting that they may be remnants from ancient rearrangements. The intact *P. falciparum* gene is *var2csa,* a var-like gene that is highly conserved between *P. falciparum* isolates^32^, involved both in cytoadherence in the placenta in primigravidae, and proposed to be a central intermediate in *var* gene switching during antigenic variation^33^. We therefore propose *var2csa* is a remnant of an ancient multigene family that has been maintained as a single complete gene in *P. falciparum,* for the dual purposes of var-switching and placental cytoadherence.

There is other evidence of retention of ancient *var* gene sequence across the subgenus. First, in Clade B we find a nearly full length *var* pseudogene that has highest similarity to *P. adleri* and *P. gaboni var* genes, within an internal *var* cluster on chromosome 4 in *P. falciparum* and *P. praefalciparum* but on the opposite strand to the other *var* genes. It is found in all *P. falciparum* isolates, but not in *P. reichenowi.* Second, in *P. gaboni* and *P. adleri,* three genes have the N-terminal DBLa/CIDRa architecture typical of Clade B genes and their domains cluster within Clade B based on similarity (Fig. 5b, larger nodes). Directly adjacent to two of these *var* genes are two *rif* pseudogenes that also show greatest similarity to those from Clade B. Last, we find a further nine *rif* pseudogenes of Clade A parasites that cluster with Clade B *rif* genes (Fig. 4). If these observations reflect retention of ancient copies, their high sequence conservation suggests that they are under extremely unusual selection pressure. Alternatively, they may represent relics of gene transfer between species that occurred after the Clade A/B split.

**Figure 5.**
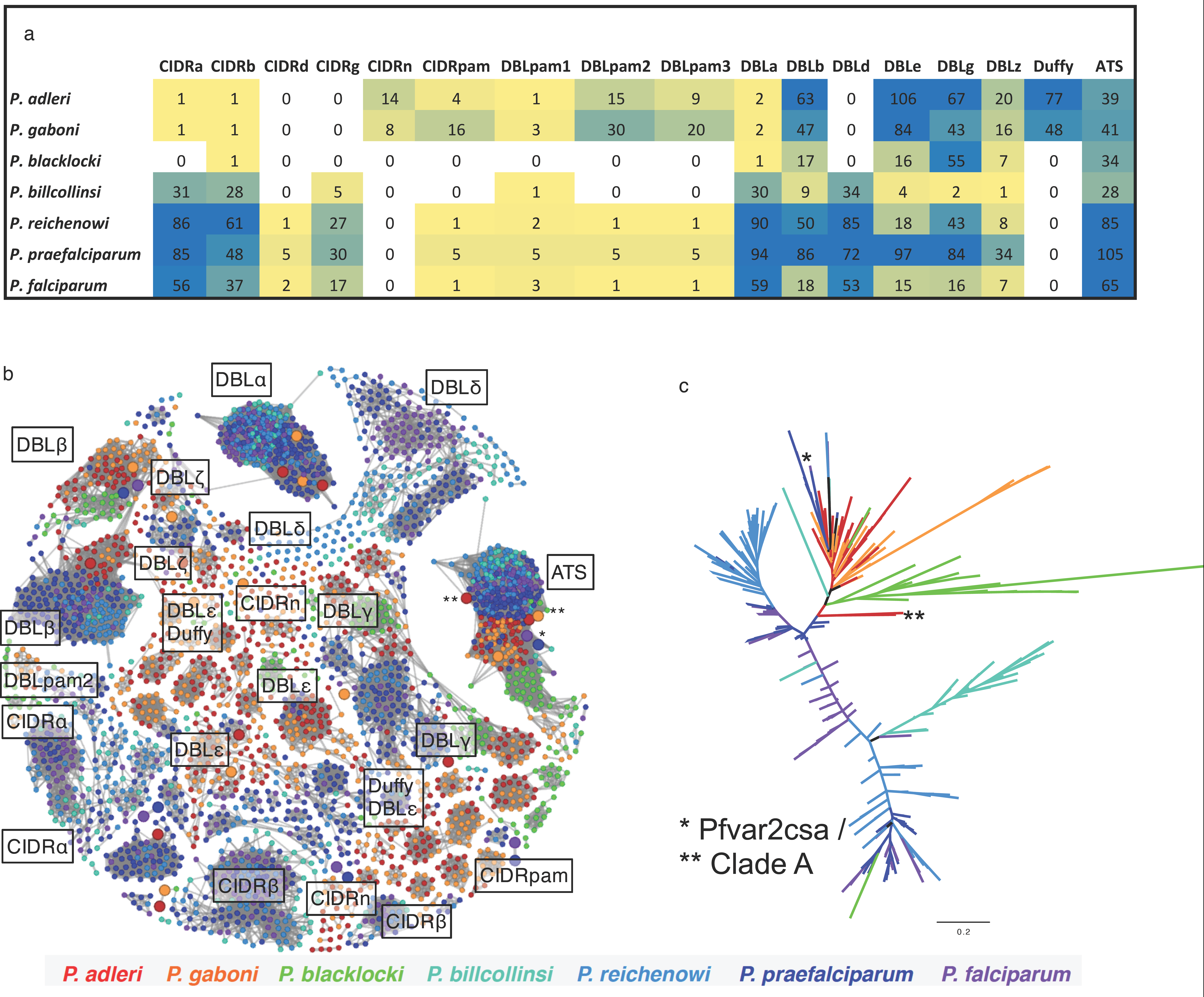
Evolution of *var* gene domains in the *Laverania*. **(a)** Heatmap of numbers of *var* gene domains in each *Laverania* species. Duffy represents regions closest to the Pfam Duffy binding domain. CIDRn is a new domain discovered in this study in Clade A. Only domains from *var* genes longer than 2.5 kb were considered. Heat map colours blue-yellow-white indicate decreasing copy numbers. (**b**) Graphical representation of similarity between domains, using domains from *var* genes longer than 2.5kb. Domains are coloured by species and clustered by a minimum BLAST cut√off of 45% global identity. Larger circles denote *var* genes in the opposite orientation. (**c**) Maximum likelihood trees of the Acidic Terminal Sequence (ATS). Apparent ATS sequences from clade A that cluster with clade B are indicated (**).

### Conclusion

We have produced high quality genomes and used mutation rates and generation times, covering the full range of most recent estimates, to calculate the date of speciation for all known members of the *Laverania,* with only a small margin of error. In our analysis, we have shown that the successful infection of humans by *P. falciparum* occurred quite recently and involved numerous parasites rather than a single one as previously proposed. After the establishment in its new host, the parasite population went through a bottleneck around 5,000 years ago during the period of rapid human population expansion due to farming (Fig. 1b). We summarise the major genomic events during the evolution of the *Laverania* in Fig. 6.

**Figure 6.**
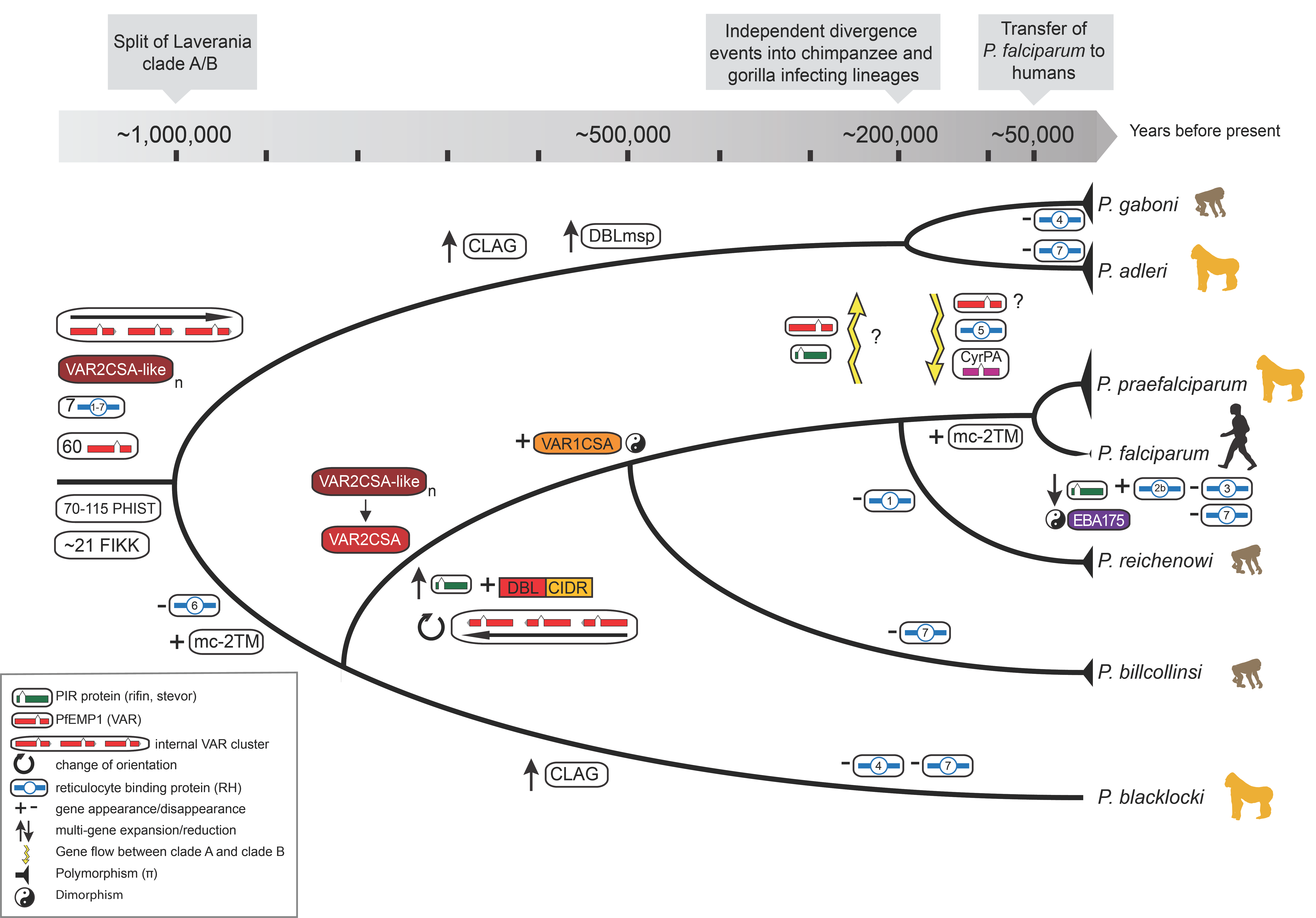
Overview of the genomic evolution of the *Laverania* subgenus. The values of polymorphism (n) within the species are indicated by triangles of different size at the end of the tree branches, as well the bottleneck in *P. falciparum* (constricted branch width), ∼ 5,000 years ago. Also shown are the gene transfers that occurred between certain Clade A and B species and the huge genomic differences that accumulated in Clade B after the divergence with *P. blacklocki.*

As a result of our analyses we propose the following series of events for the emergence of *P. falciparum* as a major human pathogen. First, the crucial lateral transfer event of the *rh5* locus between Clade A and B parasites may also have involved *var* and *rifin* genes in other parts of the genome that, because of their orientation on the opposite strand, were not lost during later recombination. Next, facilitatory mutations are likely to have occurred in *rh5* that in the first instance allowed invasion of both gorilla and human red cells. Modern humans emerged more than 300,000 years ago^34^ and existed as small isolated populations^12^. Our evidence suggests that *P. falciparum* and *P. praefalciparum* started to diverge around 40,000-60,000 years ago. In the following 40,000 years with low population densities in humans and gorillas there would have not been high selection pressure to optimise infectivity in either the hosts or vectors, enabling at least some movement of parasites between hosts. We find evidence for gene flow between lineages throughout this period. The expansion of the human population with the advent of farming likely led to strong evolutionary pressure for mosquito species (specifically *An. gambiae)* to feed primarily on humans^35^. Therefore, the existing human infective *(P. falciparum)* genotypes would be selected for human and appropriate vector success and the fittest would rapidly expand. Subsequent rapid accumulation of mutations that favoured growth in humans, and in the anthropophilic vectors such as *An. gambiae,* are likely to have occurred to increase human-specific reproductive success. The resulting specific parasite genotypes that expanded (and appeared as an emergence from a bottleneck), would have had a much lower probability of a direct transfer back to apes. With experiments on gorillas and chimpanzees not possible it will be difficult directly to prove the precise combination of different alleles that allowed the emergence of *P. falciparum.* However, existing data suggest that the genes that we have implicated in this process are expressed throughout the life cycle but that half remain uncharacterised (www.genedb.org, www.plasmodb.org), opening up new opportunities for future studies on host specificity and host adaptation in *Plasmodium.*

## Online Methods

### Sample collection

All but two infected blood samples from chimpanzees *(Pan troglodytes troglodytes)* and gorillas *(Gorilla gorilla gorilla)* were obtained from the sanctuary “Parc de La Lékédi”, Bakoumba (Haut-Ogooué, Gabon), during routine sanitary controls of the animals. This park holds various primate species, including gorillas, chimpanzees and monkeys (*Cercopithecinae*), that have been orphaned due to bushmeat-poaching activities and have been confiscated by the Gabonese Government, quarantined at the Centre International de Recherches Médicales de Franceville (CIRMF, Gabon) and finally released into semi-free ranging enclosures in the sanctuary. Every six months, chimpanzees (12 individuals) and gorillas (2 individuals) are anesthetized for medical check up. Blood samples were collected from the animals during sanitary controls (July 2011, September 2012, May 2013 and December 2013). Two additional infected blood samples were obtained from gorilla orphans (GG05, GG06) seized by the Gabonese government in 2011 and 2013 and sent to the CIRMF for a quarantine before being released in a sanctuary. All animal work was conducted according to relevant national and international guidelines. From each animal, 15 ml of whole blood were collected in EDTA tubes. For all samples but three, white blood cell depletion was performed on 10 ml of the freshly collected samples using cellulose columns as described in ^36^. Remaining blood was subsequently used for DNA extraction and detection of *Plasmodium* infections as described in Ollomo et al^3^. Overall, 15 blood samples from 7 chimpanzees and 4 gorillas were found to contain the *Laverania* samples used in the present study (Table 1).

### Sample preparation

Three methods were used for DNA amplification prior to sequencing (Supplementary Table 1). For all but one sample, whole genome amplification (WGA) was performed with a REPLI-g Mini Kit (Qiagen) following a modified protocol^37^ to enrich genomic DNA. The genome of *P. blacklocki* was generated using selective WGA (sWGA) as indicated in^38^ using 20 primers, followed by a WGA. Finally, for the PprfG03 (a *P. praefalciparum* isolate) and PadlG02 (a *P. adleri* isolate) samples, we used a cell sorting approach^39^.

### Sample sequencing

All samples were first sequenced with Illumina. Amplification-free Illumina libraries of 400-600 bp were prepared from the enriched genomic DNA^40^ and run on MiSeq and HiSeq 2000 (v3 chemistry) Illumina machines.

After the Illumina sequencing, six samples with a combination of the least number of multiple infections (see below) and the lowest level of host contamination were chosen for long read sequencing, using Pacific Biosciences (PacBio). The DNA of the samples (after WGA) was size-selected to 8 kb and sequenced with the C3/P5 chemistry. The number of SMRT cells (Pacific Bioscience sequencing runs) used varied between samples (Supplementary Table 1).

### Genome assembly, genome QC, split of infection & annotation

#### Determination of multiple infections

To initially quantify multiple infections and so allow samples to be selected for PacBio sequencing from those comprising a low number of species, Illumina reads from each sample were mapped against a concatenation of all available *Cox 3* and *CytB* genes of the *Laverania* from NCBI, using SNP-o-Matic^41^ (parameter chop=5) to position reads only where they aligned perfectly. SNP-o-Matic returns all the positions of repetitive mapping reads. This output allowed us to count the read depth of these two genes across all species and therefore determine the number and relative amount of different malaria species per sample.

#### Whole genome amplification (WGA) bias

The uneven coverage that resulted from WGA bias, host contamination and multiple infections presented a challenge for sequence assembly. To overcome the bias and the host contamination, each DNA sample was sequenced deeper than normally necessary. Lower coverage of the subtelomeres was obtained for the sWGA sample *(P. blacklocki)* meaning that the subtelomeres in that assembly were not as complete as those in the assemblies for other species.

#### Long reads (Pacific Bioscience) assemblies

Six reference genomes were assembled using HGAP^42^, with different settings for the genome size parameter, ranging from 23 Mb *(P. reichenowi)* to 72 Mb *(P. billcollinsi).* This parameter encodes how many long reads are corrected for use in the assembly and depends on the host contamination and the amount of different isolates in the samples. The obtained contigs from HGAP were ordered with ABACAS^43^ against a *P. falciparum* 3D7 reference that has no subtelomeric regions. Assembly errors and WGA artefacts were manually corrected using ACT^44^. After this step, three iterations of ICORN2^45^ were run, followed by another ABACAS step, allowing overlapping contigs to be merged (parameter: ABA_CHECK_OVERLAP=1). For the PrG01, PgabG01 and PadlG01 assembly, we also ran PBjelly to close some of the sequencing gaps^46^.

#### Host decontamination

To detect and remove sequence data derived from host DNA, contigs were compared with the chimpanzee or gorilla genomes using BLAST. Contigs were considered as host contamination if more than 50% of their BLAST hits had higher than 95% identity to any of the great ape genomes. Unordered contigs with a GC content >32% were searched against the non-redundant nucleotide database, to detect and remove further contaminants.

#### Resolving multiple infections

The first assembled genome was a single *P. reichenowi* infection, PrG01. We detected low levels of *P. vivax-like* and virus contamination (TT virus, AB038624.1), which were excluded. For quality control, the assembly was compared against the existing PrCDC^9^ reference genome. The number of *Plasmodium* interspersed repeats (PIRs) was similar, and there were no breaks in synteny. There were however significantly fewer sequencing gaps and 17 Rep20 regions could be found (a known repeat close to the telomeres in *P. falciparum)*. Thus, the assembly of PacBio data (PrG01; Supplementary Table 2) appears to be of higher quality than the existing *P. reichenowi* PrCDC reference.

The *P. adleri* sample comprised a single infection. Because a large number of cycles of amplification were used, a greater number of SMRT cells were sequenced (Supplementary Table 1) to overcome the problem of uneven coverage resulting in under-represented regions. An estimated genome size of 60 Mb was chosen for the HGAP analysis to ensure that all regions were covered.

PgabG01 was a *P. gaboni* isolate with a *P. vivax-like* co-infection. To detect contigs of *P. vivax,* unordered contigs (those that could not be placed against Pf3D7 using ABACAS) were searched against the protein sequences of *P. falciparum* 3D7 and the *P. vivax* PvP01 reference genome using TBLASTx. For each contig, the relative number of genes hitting against the two genomes was used to assign it to *P. gaboni* or *P. vivax.* In most cases, all genes for a given contig consistently hit only one genome so that the attribution to either species was clear. Overall, 14 Mb of *P. vivax-like* sequences were obtained that will be described elsewhere.

The *P. billcollinsi* genome (PbilcG01) was obtained from a co-infection with a *P. gaboni* genome (PgabG02). Rather than ordering the contigs just against Pf3D7 with ABACAS, contigs were ordered against a combined reference comprising *P. gaboni* (PgabG01) and the Pf3D7 (parameters: overlap 500 bp, identity 90%). The species designation of contigs was confirmed with a TBLASTx searches of annotated genes against a combination of the proteomes of PgabG01 and PrCDC. For subtelomeric gene families, contigs were attributed to species if the hit was significant for one species, not the other. Some of the contigs could not be attributed unambiguously and were discarded. Due to sequencing gaps, some of the core genes are missing from the final assembly.

The sample used to produce the *P. praefalciparum* genome (PprfG01) had a high level of host contamination, a low level of co-infection with *P. adleri* and contained two distinct *P. praefalciparum* genotypes. For the core genome, we used iCORN to identify and utilise the dominant genotype at each position. Where it was not possible to phase the genotypes, due to a lack of variation, we assumed that they were identical. In the subtelomeres however, it was possible to distinguish but not phase the two *P. praefalciparum* genotypes resulting in approximately twice the number of *var* genes as seen in *P. falciparum.* Due to contamination of construction vectors *(E. coli)* and host, 29 SMRT cells were sequenced and the HAGP parameter for the assembly size was set to 60 Mb. Contigs were screened against *P. adleri* and *P. falciparum* to exclude a *P. adleri* co-infection. All of the contigs that had a *P. falciparum* BLAST hit or had no clear hit (such as those containing species-specific gene families) were attributed to the *P. praefalciparum* assembly. Last, all samples (Supplementary Table 1) including five *P. falciparum* genomes were mapped against the Pf3D7, *P. praefalciparum and P. adleri* assemblies. Contigs were excluded where more normalized hits to the three *P. adleri* samples were found than to one of the two other *P. praefalciparum* samples. Similarly, this method was used to eliminate the remote possibility that any of the contigs in the *P. praefalciparum* assembly were in fact derived from *P. falciparum* co-infection.

The *P. blacklocki* sample was from a single infection. Due to sWGA, the PacBio sequence data covered regions not covered by Illumina but due to the bias of the primers, the subtelomeres were not covered fully. However, the internal *var* gene clusters are all assembled. Some of the core genes from this species are also missing.

#### Annotation

The genomes were annotated as described in^47^. In short, the annotation of *P. falciparum* (version July 2015) was transferred with RATT^48^ and new gene models were called with Augustus^49^. Obvious structural errors in core genes were manually corrected in Artemis^50^.

### Mapping - generation of further samples

To generate the gene sequence for different samples, Illumina reads were mapped against a set of reference genomes using BWA^51^ and default parameters. For the gorilla samples, we mapped against the combined PacBio reference genomes of *P. adleri, P. blacklocki* and *P. praefalciparum* and for the chimpanzee samples, the combined references of *P. gaboni* (PgabG01), *P. billcollinsi* and *P. reichenowi* (PrG01). SNPs with Phred score ≧ 100 were called using GATK UnifiedGenotyper^52^ v2.0.35 (parameters: −pnrm POOL −ploidy 2 −glm POOLBOTH). From these SNP calls we constructed the new gene set, masking regions in genes with less than 10x coverage of ‘properly’ (correct distance and orientation) mapped paired reads. To generate the sequences of the other 13 isolates, homozygous SNP calls were obtained (consensus program from bcftools-1.2^53^). We quality controlled the SNP calling by regenerating PrCDC and PgabG02 gene set from PrG01 and PgabG01, respectively and confirmed that they were placed with nearly no differences in a phylogenetic tree.

### Orthologous group determination and alignment

Orthologous groups were identified using OrthoMCL v1.4^54^ across: (i) the seven core *Laverania* genomes; (ii) the seven core genomes, the *Laverania* isolates PgabG02, PrCDC and *P. falciparum* IT, as well as two outgroup genomes *Plasmodium vivax* Sal1 and *Plasmodium knowlesi* strain H; and (iii) just Pf3D7, PprfG01 and PrG01. *P. praefalciparum* II was excluded due to its partial genome. From these groups, different complete sets of 1:1 orthologues were extracted:

1. “Lav12sp” set of 3,369 orthologues across the seven core *Laverania* species, the PrCDC and *P. falciparum* IT isolates, *P. vivax* and *P. knowlesi*
2. “Lav25st” set of 424 1:1 orthologues from across the 25 *Laverania* isolates, including the previously published *P. reichenowi* CDC and five *P. falciparum* isolates (3D7, IT, DD2, HB3 and 7G8^9^).
3. “Lav7sp” set of 4,350 orthologues from across the seven *Laverania* reference genomes
4. “Lav15st” set of 3,808 orthologues, with two representative sequences per species, excluding *P. blacklocki* and *P. praefalciparum.*
5. “Lav3sp” set of 4,826 1:1 orthologues across all the *P. reichenowi, P. praefalciparum* and *P. falciparum* isolates

The first two sets were used to reconstruct the species tree, the third one for the comparative genomic analyses (introgression, convergence and gene family evolution), the fourth one for the analyses of within species polymorphism and the fifth one for the analysis of *P. falciparum* adaptive evolution.

To reduce the rate of false positives in the evolutionary analyses due to misalignments (e.g.^55^), codon-based multiple alignments were performed using PRANK^56,57^ with the −codon and +F options, as it was shown to outperform other programs in the context of the detection of positive selection^58,59^. Prior to aligning codons, low complexity regions were excluded in the nucleotide sequences using dustmasker^60^ and in amino acid sequences using segmasker^61^ from NCBI-BLAST. Poorly aligned regions were excluded using Gblocks^62^, with default settings.

### Analysis of interspecific gene flow, introgression or gene transfer

#### Species-tree inference

Two ML trees were performed using RAxMLv8.1.20^63^ to illustrate the phylogenetic relationships between the *Laverania* species and genotypes studied here using the “Lav12sp” and the “Lav25st” set of orthologues. For each tree, multiple nucleotide alignments of each orthologous group were conducted as described above. Trees were then constructed from the concatenated alignments of the “Lav12sp” set of orthologues for the species tree and the “Lav25st” set for the strain tree using RAxML and the following options “-m GTRGAMMA –f a -# 100”. Trees were rooted afterwards using *P. vivax* and *P. knowlesi* for the species tree and the *P. adleri/P. gaboni* clade for the genotype tree.

#### Tree topology test

Interspecific gene flow was investigated by testing congruence between each gene tree topology and the species tree topology. We performed the Shimodaira-Hasegawa test (SH test^64^) using RAxMLv8.1.20 to test whether the phylogenetic tree for each gene significantly differed from the *Laverania* species tree. Topology tests were based on multiple nucleotide alignments of the 4,350 “Lav7sp” set of orthologues. For each coding sequence, RAxML was called with the options “-m GTRGAMMA –fh”.

### Convergent evolution analyses

#### Genome-wide test of convergent evolution

Convergent substitutions can occur by chance and the number of random convergent substitutions between two lineages is correlated with the number of divergent substitutions observed in these two lineages^65,66^. Excess of convergent substitutions in specific branch pairs can thus be identified by analyzing the correlation between the number of convergent and divergent substitutions between all the branch pairs in a phylogeny using orthogonal regression, and looking for outlier branch pairs: branch pairs with a high positive residual show an excess of convergent substitutions relatively to the number of divergent substitutions^65^. We used the software Grand-Convergence (available at https://github.com/dekoning-lab/grand-conv) to estimate for each chromosome the numbers of divergent and convergent substitutions between all branch pairs in the *Laverania* tree and investigate whether branch pairs including *Laverania* species infecting the same host species (gorilla or chimpanzee) presented an excess of convergence. Analyses were performed under different models of amino-acid evolution: LG, WAG, JONES and DAYHOFF.

#### Gene-based detection of convergent evolution throughout the Laverania

For each orthologue of the “Lav7sp” set, the number and percentage of fixed amino acid differences between parasites infecting the same host were calculated, *i.e.* the number of positions showing the same amino acid within a host species but different amino acid between host species. Alignments of all the available sequences (“Lav15st”) from all the sequenced isolates were then used to determine what number of host-specific differences were fixed within each host and each species. To evaluate whether the observed number of host-specific fixed differences in an alignment can be attributed to neutral evolution/purifying selection alone (with no positive selection), we used a simulation-based approach. For each coding sequence, 1,000 sequences of the same size were simulated, evolving along the same tree with the same specified branch lengths, substitution model, codon frequencies and omega (*d*_N_/*d*_S_), using the program Evolver from PAML v4.8a^67^. The program Codeml from PAML v4.8a^67^ was first used to estimate the tree, the codon frequencies and the average omega values for each of the coding sequences with fixed amino acid differences. For each simulated dataset, the number of fixed amino-acid differences between the parasites infecting a same host was estimated. The probability of observing *n* fixed differences was then computed as the proportion of the simulated dataset of 1000 sequences that showed at least the same number of fixed differences as observed in the real data.

### Tests for positive selection

#### Branch site tests

To search for genes that have been subjected to positive selection in the *P. falciparum* lineage alone after the divergence from *P. praefalciparum*, we used an updated branch site test ^68^ implemented in PAML v4.4c ^67^. This test detects sites that have undergone positive selection in a specific branch of the phylogenetic tree (foreground branch). The “Lav3sp” set of 4,826 orthologous groups between *P. reichenowi, P. praefalciparum I* and *P. falciparum* was used for the test. *d*_N_/*d*_S_ ratio estimates per branch and gene were obtained using Codeml (PAML v4.4c) with a *free-ratio* model of evolution. This identified 139 genes with a significant signal of positive selection in *P. falciparum* only.

Using the “Lav7sp” dataset, a branch site test was also applied for each gene (Figure 2), comparing the sequence from an individual species to the inferred sequence of its closest branch point on the tree (e.g. *P. falciparum* compared with the sequence at the *P. falciparum*– *P. praefalciparum* split).

#### McDonald–Kreitman (MK) tests

Selection in *P. falciparum* was also tested using McDonald–Kreitman (MK) tests ^69^ to compare the polymorphism within species to the divergence between species, using *P. praefalciparum* as the outgroup. Analyses were performed using the 4,826 “Lav3sp” set of orthologues. MK tests were performed as described before^9^. Thirty-five genes had an MK ratio considerably higher than 1.

### Gene Ontology enrichment analyses

Analysis of Gene Ontology (GO) term-enrichment was performed in R, using TopGO^70^ with default parameters. GO annotations from GeneDB were used but with unreviewed automated annotations excluded.

### Gene family analyses

To estimate the differential abundance of gene families across species, the Gene products and the Pfam domains were counted and analysed by the variance of the occurrence. Unless otherwise stated, trees were constructed using PhyML^71^ (default parameters) or RAxML^63^ (model estimated) from alignments generated with Muscle^72^ and trimmed with Gblocks^62^ in Seaview^73^ with default values. Many of the findings were confirmed manually through ACT and bamview^50^. The analysis of the *var* genes was performed on *var* genes larger than 2.5kb. Domains were called with the HMMer models from varDom^74^. Distance matrices were generated based on BLASTp scores, without filtering low complexity regions. Representation was done in R through the heatmap.2 program from gplot (see also Supplementary Note 3).

#### Allelic dimorphisms

For the analysis of dimorphism in *msp,* all sequences available for the *Laverania* were downloaded from Uniprot^75^. Data were subsampled to obtain a similar number of sequences for each group. Phylogenetic trees were constructed with PhyML^71^, using default parameters and drawn in Figtree. The *eba-175* alignment was visualized with Jalview^76^.

### Divergence Dating

Alignments of the *Laverania* included intergenic regions where possible. Assuming 402–681 mitotic events per year (Supplementary Note 1) and a mutation rate of 3.78E-10 for 4 mitotic events^77,78^(mutation rate from latter paper was taken from Pf3D7 line without drugs), equivalent to around 0.9–1.5 mutations per genome per year. Although we observed similar mutation rates in clinical samples (Supplementary Note 1), these estimates have potential errors and therefore we report ratios of divergence times in the figures that are robust to errors in these parameters. For coalescence based estimates of speciation times, G-Phocs^79^ was used and multiple sequentially Markovian coalescent (MSMC) on segregating sites^80^ was used to estimate the *P. falciparum* bottleneck.

## Author Contributions

TDO, BO, FR, CN, MB, FP designed the study. CA, APO, LB, EW, BN, ND, CP, PD, VR, FP collected and assessed samples. CA performed the WGA and cell sorting on one sample. SO performed the WGA on the samples; MS organised the sequencing. TDO did assembly and annotation. UB did manual gene curation; AG, FP performed the evolutionary analyses on core genomes. TDO, CN, MB performed the analyses of gene families and dimorphisms. TC performed the dating analyses. TDO, AG, CN, MB, FP wrote the manuscript. All authors read and approved the paper.

## Acknowledgments

This work was funded by ANR ORIGIN JCJC 2012, LMI ZOFAC, CNRS, CIRMF, IRD and the Wellcome Trust (grants WT 098051 and WT 206194 to the Sanger Institute, 104792/Z/14/Z to CN). TC holds a MRC DTP Studentship. We thank Gavin Rutledge for performing the sWGA and Julian Rayner and Francisco J. Ayala for helpful discussion. We thank the PlasmoDB team for promptly making these data available.

## Data availability

All sequences have been submitted to the European Nucleotide Archive. The accession numbers of the raw reads, and assembly data can be found in Supplementary Table 8. The genomes are being submitted to EBI, project ID PRJEB13584. The genomes are available from plasmodb.org and from ftp://ftp.sanger.ac.uk/pub/project/pathogens/Plasmodium/Laverania/.

## Competing financial interests

None

## Computer code

Custom computer code is available on request.

